# ROOT PENETRATION INDEX 3, a major quantitative trait locus (QTL) associated with root system penetrability in Arabidopsis

**DOI:** 10.1101/2021.12.23.473296

**Authors:** Elohim Bello Bello, Thelma Y. Rico Cambron, Rubén Rellán Álvarez, Luis Herrera Estrella

**Affiliations:** Unidad de Genómica Avanzada/LANGEBIO, Centro de Investigación y de Estudios Avanzados, Irapuato, México; Department of Molecular and Structural Biochemistry, North Carolina State University, Raleigh, NC, USA; Institute of Genomics for Crop Abiotic Stress Tolerance, Department of Plant and Soil Science, Texas Tech University, Lubbock, TX 79409, USA

**Keywords:** Arabidopsis accessions, mechanical impedance, natural variation, QTL mapping, root morphology, root system penetrability

## Abstract

Soil mechanical impedance precludes root penetration, confining root system development to shallow soil horizons where mobile nutrients are scarce. Using a two-phase-agar system, we characterized *Arabidopsis thaliana* responses to low and high mechanical impedance at three root penetration stages. We found that seedlings whose roots fail to penetrate agar barriers show drastic changes in shoot and root morphology, while those capable of penetrating have only minor morphological effects. The assessment of 21 Arabidopsis accessions revealed that primary root penetrability (PRP) varies widely among accessions. To search for quantitative trait loci (QTLs) associated to root system penetrability, we evaluated a recombinant inbred population (RIL) derived from Landsberg erecta (Ler-0, with a high PRP) and Shahdara (Sha, with a low PRP) accessions. QTL analysis revealed a major-effect QTL localized in chromosome 3 (*q-RPI3*), which accounted for 29.98% (LOD = 8.82) of the total phenotypic variation. Employing an introgression line (IL-321), with a homozygous *q-RPI3* region from Sha in the Ler-0 genetic background, we demonstrated that *q-RPI3* plays a crucial role in root penetrability. This multiscale study revels new insights into root plasticity during the penetration process in hard agar layers, natural variation and genetic architecture behind primary root penetrability in Arabidopsis.

**Highlight:** We found a wide natural variation in the capacity of Arabidopsis accessions to penetrate hard agar layers. Using a Ler-0 x Sha recombinant inbred population, a major-effect QTL (*q-RPI3*) strongly associated with root penetrability of compact agar layers was identified.

## Introduction

Plant roots growing on compacted soils are usually exposed to a multi-stress environment, experiencing high mechanical impedance, low oxygen levels, water stress and nutrient deprivation (Colombi and Keller, 2019; Correa *et al*., 2019; Wang *et al*., 2019). To cope with edaphic complexity of soil compaction, roots have evolved different adaptive mechanisms, which include a broad range of anatomical, morpho-physiological and molecular responses (Monshausen *et al*., 2009; Lipiec *et al*., 2012; Okamoto *et al*., 2021). The impact of soil impedance on root penetration has been described in both monocot and dicot plant species (Iijima *et al*., 1991; Tracy *et al*., 2012; Grzesiak *et al*., 2013). Despite differences in root architectures between monocots, characterized by a fibrous root system, and dicots, generally showing taproot systems, high soil mechanical impedance (>2 MPa) severely affects root system development across all plant types. Compacted soils impact root system architecture by reducing total root area, root length, rooting depth and lateral root number; thus, harming root system foraging capacity and function (Bengough *et al*., 2011; Potocka and Szymanowska, 2018).

In compacted soils, root systems must optimize their growth strategies by an adaptive plasticity to either avoid impenetrable obstacles or break through hard soil layers (Jin *et al*., 2013). In this context, it has been suggested that touch-dependent gravitropic and thigmotropic responses, which include sundry root growth patterns such as root tortuosity/circumnutation function (Taylor *et al*., 2021), buckling (Silverberg *et al*., 2012), skewing (Roy and Bassham, 2017), coiling/curling (Lourenco *et al*., 2015) and waving (Tan *et al*., 2015), could facilitate root penetration into hard surfaces. At anatomical and morphological levels, a plethora of root specific traits have been reported to be associated with root penetrability such as cortical thickness, stele diameter (Chimungu *et al*., 2015), root hair density (Bengough *et al*., 2016), mucilage production (Iijima *et al*., 2004), sloughing of root cap cells (Bengough and McKenzie, 1997) and root tip geometry (Colombi *et al*., 2017; Roué *et al*., 2020). In maize (*Zea mays*) and wheat (*Triticum aestivum*), root genotypes with multiseriate cortical sclerenchyma showed greater lignin concentration and bending strength, which in turn, improve their penetration ability in compacted soils (Schneider *et al*., 2021).

The cross-talk between ethylene and auxin signal transduction pathways has been reported to be an important determinant for root penetration capacity in tomato (*Solanum lycopersicum*) (Santisree *et al*., 2011). Diverse studies on root adaptations to impenetrable barriers or touch stimuli support the notion that changes in root morphology are triggered by ethylene-induced auxin biosynthesis and redistribution (Okamoto *et al*., 2008; Jacobsen *et al*., 2021). Moreover, ethylene has been strongly implicated not only in the root system responses to mechanical impedance but also as a key signal to warn roots of the soil compacted zones (Pandey *et al*., 2021). Interestingly, it has been suggested that ethylene insensitive root systems have better chances of penetrating into compacted soils as they can maintain a steady growth through channels and pores present in the subsoil (Pandey *et al*., 2021; Vanhees *et al*., 2021). More recently a metabolome, hormonal and gene expression analysis of tomato seedlings grown under different agarose hardness levels identified abscisic acid (ABA), trehalose/SnRK and E2Fa as major modulators of mechanical stress-induced growth inhibition (Kumari *et al*., 2021). These inter-related studies denote that, like other specific plant responses to mechanical impedance, the root penetration process appears to be controlled by a complex genetic–hormonal regulatory network.

Analyses of mechanosensing and signal transduction in *A. thaliana* roots provided guidelines to understand how plant roots perceive, respond and penetrate compacted layers (Hamant and Haswell, 2017). Considerable attention has been devoted to putative plasma membrane-associated mechanosensors as important components of the molecular mechanisms regulating root responses to mechanical stress. A prominent example is *mid1*-complementing activity 1 (MCA1), a mechanosensitive ion channel localized in root cells, which mediates Ca^2+^ influx in response to mechanical stimulation. The functional characterization of the *mca1* knock-out mutant revealed that its primary roots are incapable of penetrating hard agar layers, suggesting that MCA1 plays a crucial role in touch sensing in Arabidopsis roots (Nakagawa *et al*., 2007). Similar results were obtained for the receptor-like kinase FERONIA; genetic disruption of FERONIA results in defective root growth patterns under mechanical perturbation, including an increase in root skewing and a reduction in root penetration percentages in a dense medium (Shih *et al*., 2014). Roots of *pzo1 mutants (pzo1-5* and *pzo1-6)* are also defective at penetrating denser media, which implies that Arabidopsis PIEZO (PZO1) activity is required for proper root penetration of compacted environments and plays a functional role in root mechanotransduction (Mousavi *et al*., 2021). Another important but less studied aspect of root penetration capacity is the natural variation of this trait. Some studies already highlighted the inter-and intra-specific variation in the capacity of plants roots to overcome soil physical barriers. Genotypic variation in root system penetrability has been reported for rice (Yu *et al*., 1995), soybean (Bushamuka and Zobel, 1998), wheat (Whalley *et al*., 2012), cotton (Klueva *et al*., 2000) and maize (Chimungu *et al*., 2015). It has been proposed that roots of dicots species have better penetrability than monocots species due to differences in root diameter and maximum root growth pressure (Materechera *et al*., 1991). Later studies proved that dicot roots have similar axial growth pressures than monocots, implying that their growth pressure, at least in seedling stage, do not explain the differences in their penetrability (Clark and Barraclough, 1999). In monocots, differences in root penetrability were closely connected with variation in specific root anatomical traits, such as cortical traits, which are ultimately mediated by differences in hormonal responsiveness (Schneider *et al*., 2021). However, despite significant efforts to date, the genetic mechanism underlying natural variability in root penetration remains largely unknown.

In this study, we provide new insights into *A. thaliana* responses to mechanical impedance. We show that development of Arabidopsis shoot and root growth is severely affected when root penetration was impeded by an agar layer. The observed morphological alterations in impeded roots correlated with inhibition of cell elongation and changes in the expression pattern of the auxin transporter PIN2. We also revealed an ample natural variation in the root capability of Arabidopsis accessions to penetrate hard agar layers. Finally, we report the identification of a major-effect QTL (*q-RPI3*) associated root system penetrability in Arabidopsis.

## Materials and methods

### Plant materials and growth conditions

*Arabidopsis thaliana* accession Columbia-0 (Col-0) and the previously described *QC46::GUS* (Sabatini *et al*., 1999), *CycB1;1::GFP* (Colón-Carmona *et al*., 1999), *DR5::GFP* (Ulmasov *et al*., 1997), *PIN2::PIN2::GFP* (Xu and Scheres, 2005) reporter lines were from laboratory stock. Arabidopsis accessions and *nced3* T-DNA mutant (GABI_129B08) used in this study were obtained from the Arabidopsis Biological Resource Centre (ABRC). The Ler-0 x Sha RIL population was originally obtained from the Nottingham Arabidopsis Stock Center (NASC, stock no. N24481). Seeds were surface sterilized with absolute ethanol for 7 min, 20% (v/v) commercial bleach for 7 min and rinsed three times with sterile distilled water. Sterilized seeds were vernalized for 48 hours at 4 °C and grown for 12 days after sowing (das) in 0.1X Murashige and Skoog (MS) medium, pH 5.7, supplemented with 0.5% (w/v) sucrose, 3.5 mM MES (Sigma-Aldrich) and different agar concentrations (Agar, Plant TC, Micropropagation Grade, Gel strength: 1165 g/cm^2^, PhytoTechnology Laboratories, Kansas, USA). Seedlings were grown in vertically oriented plates (90°) in a Percival chamber at 22 ±1 °C, under a long-day photoperiod (16/8 h) with ∼80 μmol·m^−2^·s^−1^ photon flux density.

### Two-phase-agar system

This system was prepared based on the method reported previously, with minor modification (Nakagawa *et al*., 2007). Briefly, the bi-phase medium consisted of a thin upper layer with a standard agar concentration (1%, w/v) and a thick bottom layer with different agar concentrations (0.6-2%, w/v). For bottom agar layer, 130 ml of MS medium was poured and spread in circular plates (Star™ Dish, 150 x 15 mm, Phoenix Biomedical, Ontario, Canada). After complete solidification, the agar layer was cut off in half and one of the gel portions was extracted. The empty area was filled with 15 ml of 1% agar MS medium. Seeds were sown at 5 mm above bottom layer. In order to obtain reproducible results, it is recommended the use of the same lot of agar throughout all the experimental procedures.

### Penetrometer resistance test

The hardness test of different agar layers was carried out by penetrating horizontally placed layers with a digital penetrometer FR-5120 (QA Supplies, Virginia, USA) equipped with a 3 mm diameter cylindrical blunt-end tip. Penetrometer resistance (PMR) was calculated as the maximum penetrating force (N) required to push the penetrometer tip into agar matrix (at a depth of 1 cm) divided by the cross-sectional area of tip base (mm^2^). The penetration resistance values are expressed as MPa.

### Root penetration index

The root penetration ability into two-phase-agar system of Arabidopsis accessions and RIL population was defined as root penetration index (RPI). The RPI was calculated as Υ = P/N, where P= the number of seedling root’s that penetrated the agar, N = number of seedlings per plate. The experimental unit was a petri dish, where around 45-50 seedlings for each accession and RIL were evaluated. Phenotypic values were from three independent experiments. For the Arabidopsis accessions, Col-0 accession was used as a control, while for the RIL population, accessions Ler-0 and Sha were established as a positive and negative controls, respectively.

### Evaluation of seedling growth and root system architecture

To characterize shoot and root phenotypes, seedlings were extracted from agar layers, except for the measurement of the rooting depth. For leaf area, shoots were separated from roots and photographed using an AxioCam ICc5 camera (Carl Zeiss, Jena, Germany) coupled to an Olympus SZH-10 stereomicroscope (Olympus, Tokyo, Japan). The images were analyzed using Fiji software (version 2.0.0) (Schindelin *et al*., 2012). For the characterization of root system architecture, primary root length was measured from root tip to hypocotyl base. The rooting depth inside agar layers was defined as primary root length from the agar inter-phase base to root tip. Root diameter was measured at root differentiation zone. The root tip index was determined as the ratio of the quiescent center (QC) diameter and the length from QC to the root cap. Meristem length and cortical cell number were measured as described (Xie *et al*., 2020). The length of elongation zone was estimated as reported (Jacobsen *et al*., 2021). The root diameter and root tip measurements were determined by Q-Capture Pro7 imaging software (QImaging, Surrey, Canada) under an Olympus BX51 microscope (Olympus, Tokyo, Japan).

### GUS staining and GFP analysis

For histochemical GUS staining, seedlings were incubated at 37 °C for specified time in a GUS staining buffer (1 mM X-gluc, 100 mM sodium phosphate, pH 7.0, 10 mM EDTA, 0.5 mM K_4_Fe(CN)_6_ and 0.5 mM K_3_Fe(CN)_6_). Stained roots were optically clarified as reported (Malamy and Benfey, 1997). Images of representative roots were captured using Nomarski optics on a Leica DMR microscope (Leica Microsystems, Wetzlar, Germany). GUS intensity quantification was performed according to the method reported previously (Béziat *et al*., 2017). Relative GUS expression was calculated by normalizing intensity values to control conditions. For analysis of the GFP expression patterns, Arabidopsis roots were stained with a 10 μg/ml propidium iodide (PI) solution (Sigma-Aldrich). GFP fluorescence was visualized by helium-neon green laser at 488 nm of excitation. PI was excited at 514 nm by diode laser line. Images were taken in the LSM-800 confocal laser-scanning microscope (Carl Zeiss, Jena, Germany). Fluorescence signal was measured based on Luke Hammond’s protocol available on GitHub https://github.com/mfitzp/theolb/blob/master/imaging/measuring-cell-fluorescence-using-imagej.rst

### ABA treatments

To determine the influence of ABA on primary root penetrability in different *A. thaliana* lines, seeds were germinated in the two-phase-agar system containing different concentrations of ABA (0, 0.1, 1, and 3 μM) in the bottom agar layer. To avoid ABA diffusion and its effect on seed germination, a thin agar band was removed between the interface of upper and bottom layers. After 12 days of ABA treatment, RPI was recorded.

### QTL mapping

Prior to the QTL analysis, RPI data were square-root-transformed to better fit for normal distribution. A genetic linkage map of the Ler-0 x Sha RILs was constructed using 92 genetic markers provided by Thomas E. Juenger’s laboratory (UT, Texas, USA). QTL analysis was carried out using R/qtl package (version 1.47-9, http://www.rqtl.org) (Broman *et al*., 2003). For QTL detection, we performed a single QTL mapping for the RPI phenotype using Haley-Knott regression (1 cM step). Additional QTLs and possible QTL interactions were assessed by multiple QTL mapping using data augmentation, forward stepwise approach and mqmscan, makeqtl and fitqtl functions. LOD significance thresholds (p < 0.05) for SQM and MQM scan were determined by running 1000 permutations. QTL support intervals were obtained based on 95% Bayes credible interval. The proportion of phenotypic variance explained (PVE) by the QTL was determined using the fitqtl function in R/qtl.

### QTL validation

The Arabidopsis introgression line (IL-321) was provided by Carlos Alonso Blanco’s laboratory (CNB, Madrid, Spain). IL-321 carries a homozygous region (6 Mb) of the top arm of chromosome 3 from Sha accession in the Ler-0 genetic background. The IL-321 as well as parental lines were phenotyped using the two-phase-agar system, considering the parameters described above.

### Statistical analysis

All statistical analyses in this study were executed using R software (version 4.1.0, http://www.R-project.org). For statistical analyses of the root phenotypes among different agar treatments, ANOVAs were conducted with a significance level of p-value < 0.01. Tukey’s HSD test was subsequently used as a multiple comparison procedure using a threshold of p-value < 0.01. For RPI analysis, we performed the estimation of the true value of RPI by including confidence intervals (CI), general lineal model (GLM) and Fisher’s exact test for 2×2 contingency tables. Other statistical attributes, including Hartigan’s dip test, Spearman’s rank and Pearsońs correlation coefficients were done under R environment. Broad sense heritability (H^2^) was calculated as H^2^ = σ2g/(σ2g+σ2ε/R) with the lmer function from the lme4 package. All the phenotypic data were graphically visualized with ggplot2 package.

## Results

### Agar mechanical impedance diminishes root penetrability and rooting depth

To assess the effect of mechanical impedance on Arabidopsis primary root penetrability, we tested different agar concentrations (0.6-2%) in a two-phase-agar system, which comprises two adjoining agar layers, a thin upper layer with 1% agar concentration and a thick bottom layer, where different agar concentrations were tested (Fig. 1A, see Supplementary Fig. S1A at JXB online). Initially, we focused on evaluating the penetrometer resistance (PMR) of the agar gels (as measure of mechanical impedance), the root penetration index (RPI) that measures root penetration ability and is defined by the formula Υ = P/N, where P= the number of seedling roots that penetrated the bottom agar layer, N = total number of seedlings per plate, and the rooting depth (RD) in each agar concentration (as measure of root elongation inside agar gels). Interestingly, agar concentration and PMR were positively correlated (R^2^ = 0.99, Fig. 1B, Supplementary Table S1), indicating that the increment of the mechanical impedance varies as a function of agar concentration in the medium. In this sense, RPI and RD were negatively correlated to agar concentration (rho= −1 and R^2^ = −0.91, respectively) (Fig. 1C, D, Supplementary Table S1). A detailed analysis of the assessed root traits indicated that, when the agar concentration increased from 0.6% (PMR: 0.01 MPa) to 0.8% (PMR: 0.04 MPa), root system penetrability and rooting depth significantly decreased by as much as 50 and 20%, respectively (Fig. 1C, D). Agar concentrations above 0.8% showed a drastic decline in both root traits, until reaching a decrease of ∼98% in RPI and ∼40% in RD at the highest agar concentration (2%, Fig. 1C, D). These observations denote a clear effect of agar concentration on mechanical impedance, root system penetrability and rooting depth of Arabidopsis seedlings (Supplementary Table S2). Considering the results described above, we assigned the agar concentrations of 0.8% (PMR: 0.04 MPa and RPI: 0.46) and 1.6 % (PMR: 0.13 MPa and RPI: 0.06) as low and high mechanical impedance treatments, respectively (Supplementary Fig. S2A, B).

**Fig. 1.**
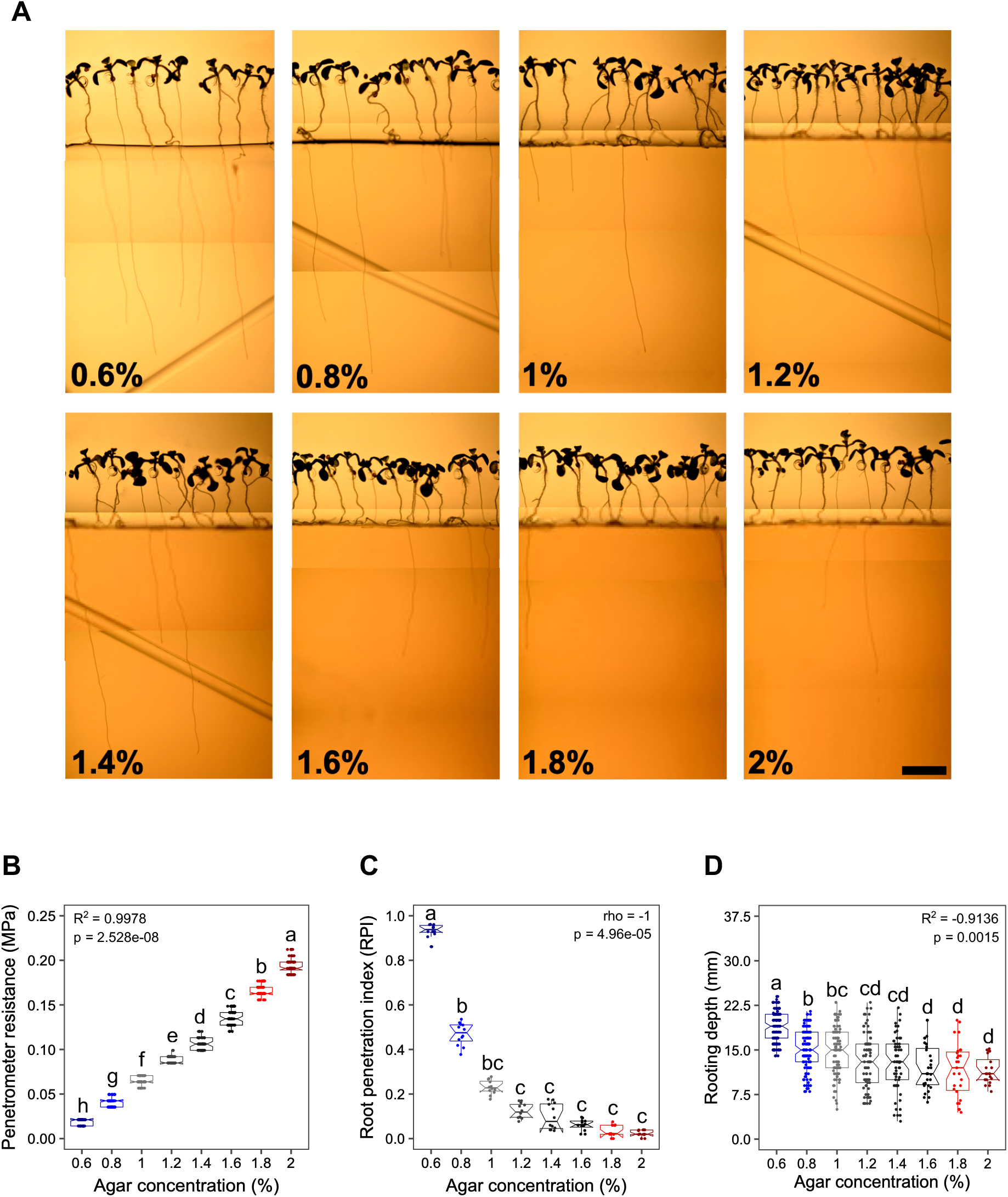
Analysis of the effect of agar concentration on mechanical impedance and root traits. (A) Representative images of Arabidopsis seedlings (Col-0 accession) growing under the two-phase culture system at different agar concentrations. Scale bar, 4 mm. Primary roots grow on the surface of the agar medium (1%) until reach the matrix of the bottom agar layer with different mechanical resistances (5-7 days). Afterwards, Arabidopsis roots penetrate the bottom layer according to agar concentration (0.6-2%). Images were reconstructed from three adjacent stacks. (B-D) Box plots of the effect of agar concentration on penetrometer resistance, root penetration index and rooting depth. Horizontal lines, medians; box limits, 25th and 75th percentiles; R^2^, Pearson’s r; rho, Spearman’s ρ; p, p-value; letters, significant differences (p=0.01, one-way ANOVA and Tukeýs HSD). Data are from three independent experiments (n= 12 plates, with 45 to 50 seedlings per plate).

### Arabidopsis seedlings exhibit distinct morphological responses during root penetration into agar layers

Having standardized the two-phase-agar system, we set out to characterize the morphological responses of 12-day-old Arabidopsis seedlings during different penetration stages in low and high mechanical impedance (Supplementary Fig. S2B, Supplementary Video S1). For the purposes of this study, we defined the Arabidopsis seedling phenotypes according to three root penetration stages (RPS) with respect to the surface of the bottom agar layer (SBAL). Stage 1 was cataloged as Arabidopsis plants whose roots did not touch the SBAL (non-disturbed roots, NDR), stage 2 as those whose roots were impeded by SBAL (impeded roots, IR) and stage 3 as those whose roots penetrated the SBAL (penetrating roots, PR) (Supplementary Fig. S1B, Supplementary Fig. S2B). In these experiments we used Arabidopsis seedlings that did not touch the SBAL (NDR) as control to characterize the effects of mechanical impedance of IR and PR roots. We first compared the impact of root penetration vs not penetration in Arabidopsis seedlings grown under the low mechanical impedance treatment. We observed significant changes in shoot and root morphology in the seedlings with IR and PR when compared to control seedlings. Seedlings which root failed to penetrate the second agar layer showed a drastic reduction in leaf area, primary root length and elongation zone, while root diameter was significantly increased (Fig. 2, Supplementary Fig. S2C). Other shoot and root traits were not affected by low mechanical stress (Supplementary Fig. S3). By contrast, seedlings which root penetrated the second agar layer showed significant reduction in leaf area, primary root length and root diameter, but much less drastic than in seedlings which root did not penetrate agar layer (Fig. 2, Supplementary Fig. S2C). No other morphological changes were observed in the rest of the assessed traits for PR roots (Supplementary Fig. S3). These results suggest that failing to break an initial root penetration barrier has a drastic effect on seedling growth, while seedlings that penetrate agar layer with low mechanical impedance only suffer a small negative impact on some specific traits. To assess seedling morphology under high mechanical impedance, we used the same comparison schema raised before, founding a similar trend in morphological changes in shoots and roots of the analyzed seedlings (Fig. 2, Supplementary Fig. S2C). We also observed that, independently of the concentration of the bottom agar layer, seedlings with penetrating roots had significantly longer roots, higher leaf area, larger elongation zones and a thinner diameter that those that failed to penetrate the SBAL (Fig. 2, Supplementary Fig. S2C). On the other hand, roots unable to penetrate showed a wavy phenotype but not those able to penetrate agar layers in both high and low impedance. Interestingly, root waving was more clearly seen in roots that remained on the surface of the lower impedance agar layer than in those on the surface of the higher impedance (Supplementary Fig. S2C). Also, roots unable to penetrate the bottom agar layer had an ectopic production of root hairs near the root tip (Fig. 2E, K). Together, these results demonstrate that mechanical impedance has an adverse effect on early stages of seedling development in *A. thaliana*. Because our root morphological results are particularly interesting in relation to root adaptive plasticity to high soil mechanical impedance, we further decided to investigate the molecular nature of root system penetration under the two-phase-agar system.

**Fig. 2.**
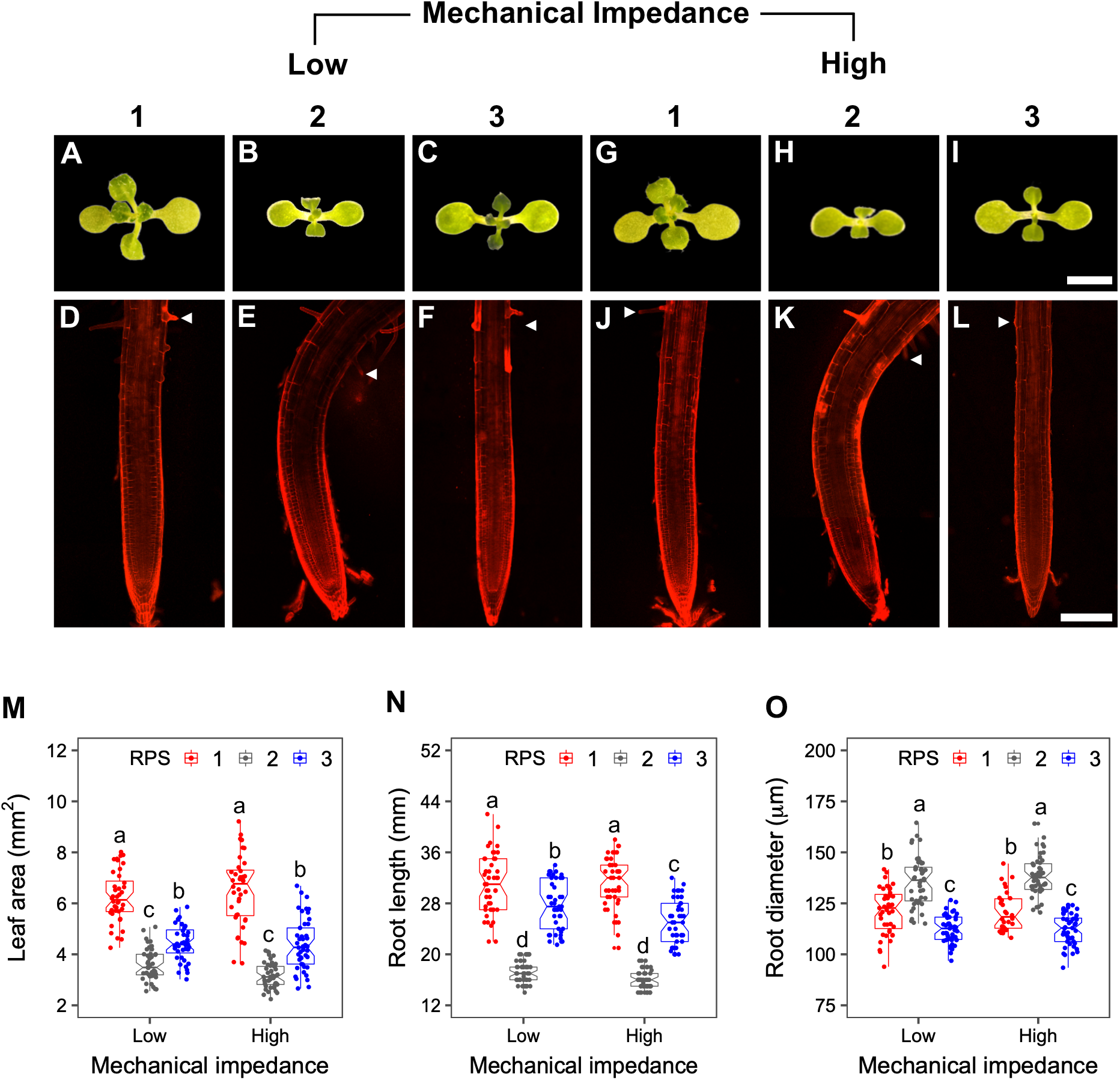
Analysis of the morphological responses of Arabidopsis seedlings grown under low and high mechanical impedance. Shoot and root morphology of 12-day-old Arabidopsis seedlings grown at three root penetration stages (RPS): before (1), during (2) and after (3) root tip touches the bottom agar layer at low (A-F) and high (G-L) mechanical impedance. Root images were fused from two adjacent stacks. White arrows, root hair localization. Scale bar, 150 μm. (M-O) Box plots show median of the tested morphological traits. Horizontal lines, medians; box limits, 25th and 75th percentiles; letters, significant differences (p=0.01, two-way ANOVA and Tukeýs HSD). Data are from three independent experiments (n= 10-12 seedlings per experiment).

### Mechanical impedance does not affect QC cell identity or cell proliferation during penetration of hard agar layers

To understand the molecular mechanisms underlying root penetrability into agar layers, we used the two-phase-agar system to examine the expression patterns of different reporter lines. In all the cases, both low and high impedance treatments were considered. Because root elongation and penetration were affected by agar impedance, we focused on analyzing the effect of mechanical stress on the quiescent center (QC) identity and meristem activity. With this aim, we used the *QC46::GUS* (QC cell identity) and *CycB1;1::GFP* (cell division) as reporter lines. GUS staining and GFP fluorescence in the tested reporter lines was similar in Arabidopsis roots that failed to penetrate and those that were able to penetrate the second agar layer in both high and low impedance treatments. These results show that stem cell activity and meristematic cell division were not affected by mechanical impedance during the root penetration process (Supplementary Fig. S4). Therefore, the drastic or slight reduction of the root length observed in the IR and PR are not due to changes in stem cell activity or meristematic cell division. These observations suggest that such root reductions found in IR and PR phenotypes could be explained mainly by alterations in cell elongation caused by continuous mechanical stress.

### Mechanical impedance alters auxin responsiveness and the expression pattern of PIN auxin transporters

In accordance with the observed results above, we asked if the IR and PR phenotypes of Arabidopsis were regulated by the plant hormone auxin, which has been extensively studied in the context of plant responses to mechanical stimuli (Rashotte *et al*., 2000; Squires and Bisgrove, 2013; Lee *et al*., 2020). To address this question, we analyzed auxin-mediated root responses through the visualization of the expression patterns of the auxin-responsive marker *DR5::GFP*. As shown in Fig. 3A and C, roots not subjected to mechanical stress expressed *DR5::GFP* in the quiescent centre and columella cells as reported previously (Ottenschläger *et al*., 2003). By contrast, we found that seedlings for which root tips were unable to penetrate agar layers (IR roots), in both mechanical treatments, exhibited an increase in GFP fluorescence intensity in the lateral root cap cells, columella cells and epidermal cells of the elongation zone. A higher GFP signal was detected in the epidermal layer of roots exposed to low impedance than those exposed to high impedance (Fig. 3A, C). When we compared seedlings with impeded and penetrated roots in the second hard agar layer with the controls, we observed a significant increase in GFP fluorescence in the quiescent centre and in three or four columella cells as compared to controls (Fig. 3A, C). Also, a significant increase in fluorescence was observed in the lateral root cap cells from the PR in high impedance conditions. Albeit subtle changes in the pattern of *DR5::GFP* expression were observed between IR and PR, no statistical differences were found in the GFP fluorescence intensity between them (Fig. 3A, C). These results suggest that auxins play an important role during the primary root penetration process into layers with different impedance levels. As changes in auxin redistribution have been reported to be involved in root movement during mechanical obstacle avoidance, we decided to analyze the PIN2 auxin efflux carrier using a marker line containing *PIN2::PIN2::GFP* gene construct. Interestingly, when seedling roots that failed to penetrate the bottom agar layer (IR phenotypes), *PIN2::PIN2::GFP* fluorescence in cortical and epidermal cells in the elongation zone of the root was significantly decreased when compared to the controls (Fig. 3B, D). However, once roots penetrate such layer (PR phenotypes), the *PIN2::PIN2::GFP* fluorescence level was restored to similar levels to the control seedlings. Although non-significant, GFP fluorescence increase in PR under high impedance was slightly higher than in control seedlings (Fig. 3B, D). These findings support that auxin activity and its redistribution dynamics across different root zones and cell types fulfils an important function during the penetrability of hard layers.

**Fig. 3.**
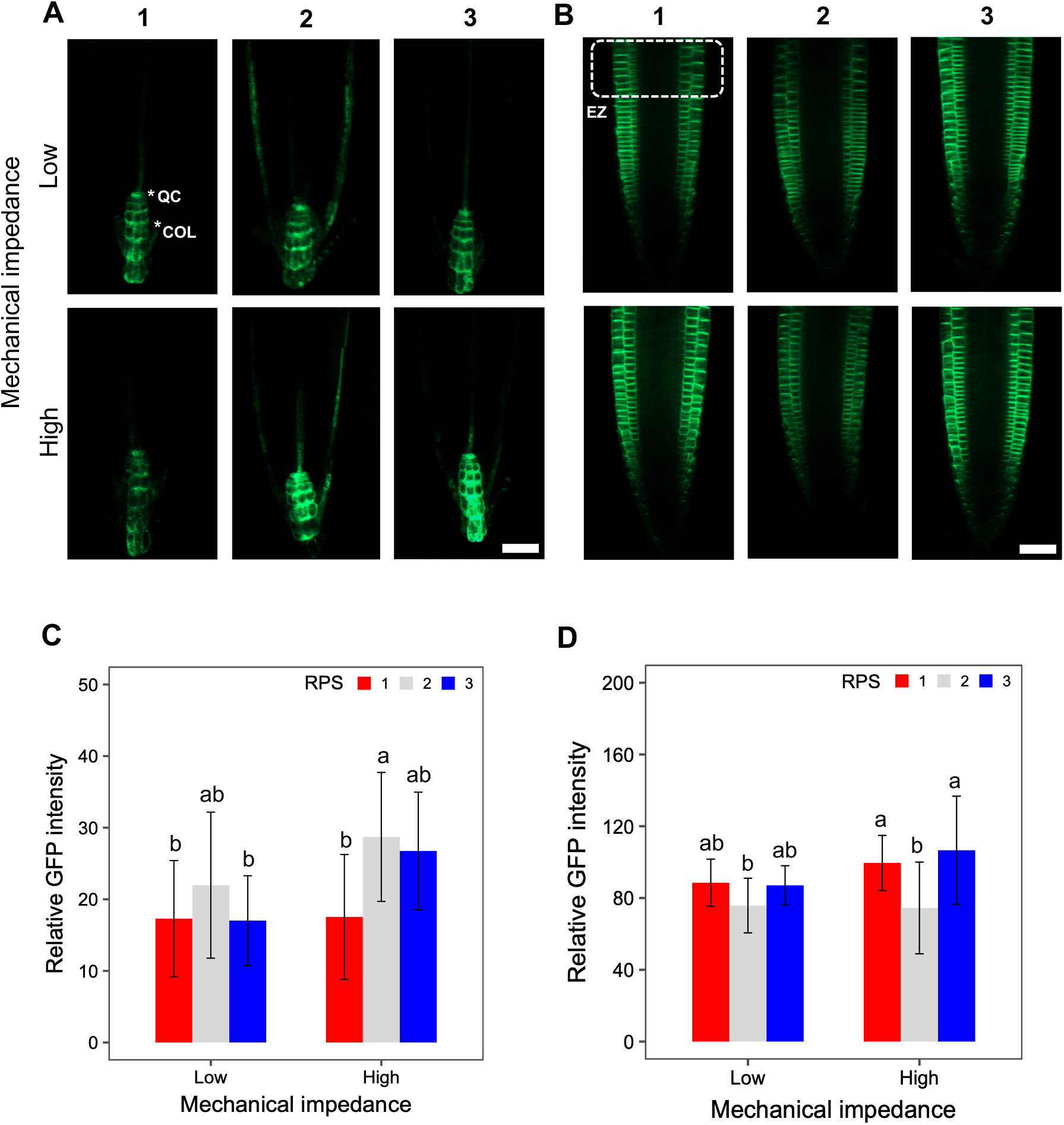
Analysis of the *DR5::GFP* and *PIN2::PIN2::GFP* reporter lines during Arabidopsis primary root penetration into agar layers. Expression patterns of *DR5::GFP* (A) and *PIN2::PIN2::GFP* (B) in Arabidopsis roots during three RPS at low and high mechanical impedance. Scale bar, 50 µm. *DR5::*GFP intensity was measured from the quiescent centre (QC) to the first three columella cells (COL), while *PIN2::PIN2::*GFP was quantified at the root elongation zone (EZ) part. Green color, GFP fluorescence. Scale bar, 50 µm. (C) and (D) Quantification of GFP fluorescence in the *DR5::GFP* and *PIN2::PIN2::GFP* reporter lines, respectively. Bar plots show the mean±standard deviation of GFP fluorescence intensity. Letters, significant differences (p=0.01, two-way ANOVA and Tukeýs HSD). Data are from three independent experiments (n=10-15 roots).

### Arabidopsis accessions have a wide natural variation in the capacity to penetrate hard agar layers

To investigate the genetic architecture of root system penetration into compacted agar media, we explored the natural variation in root penetration ability in *A. thaliana*. We used a small panel of 21 Arabidopsis accessions collected from different edaphic conditions (Supplementary Table S3). The edaphic variables at the sites of origin of the accessions cover a broad range of soil environments with very contrasting top-(0-30 cm) and subsoil (30-100 cm) physical properties, including different soil units, bulk densities and textures (Supplementary Table S3). We hypothesized that root adaptive plasticity to diverse soil physical conditions could provide insight into the genetic mechanisms underlying root penetrability into compacted soils. On that basis, we employed the two-phase-agar system with an 1% agar concentration (PMR: 0.06 MPa and RPI: 0.23) to phenotype the RPI trait among these 21 Arabidopsis accessions. Even in this small set of Arabidopsis accessions, we observed a wide natural variation in the capacity of the Arabidopsis root system to penetrate hard agar layers, with penetration indexes ranging from 0.13 to 0.51 (Fig. 4A, Supplementary Table S4). The phenotypic data analysis showed that 14 accessions exhibited a RPI phenotype similar to Col-0 (RPI=0.24, Fig. 4C) whereas accessions Uod-1, Van-0, Ler-0 and Zdr-1 showed the highest RPI values and HR-5, Sha and Ei-2 accessions the lowest RPI values (Fig. 4A, C). These findings suggest that root systems of accessions respond differently to substrate compaction. As such, the observed natural variation in the capacity of roots to overcome hard layers can be used to unravel the genetic components controlling the root penetration process.

**Fig. 4.**
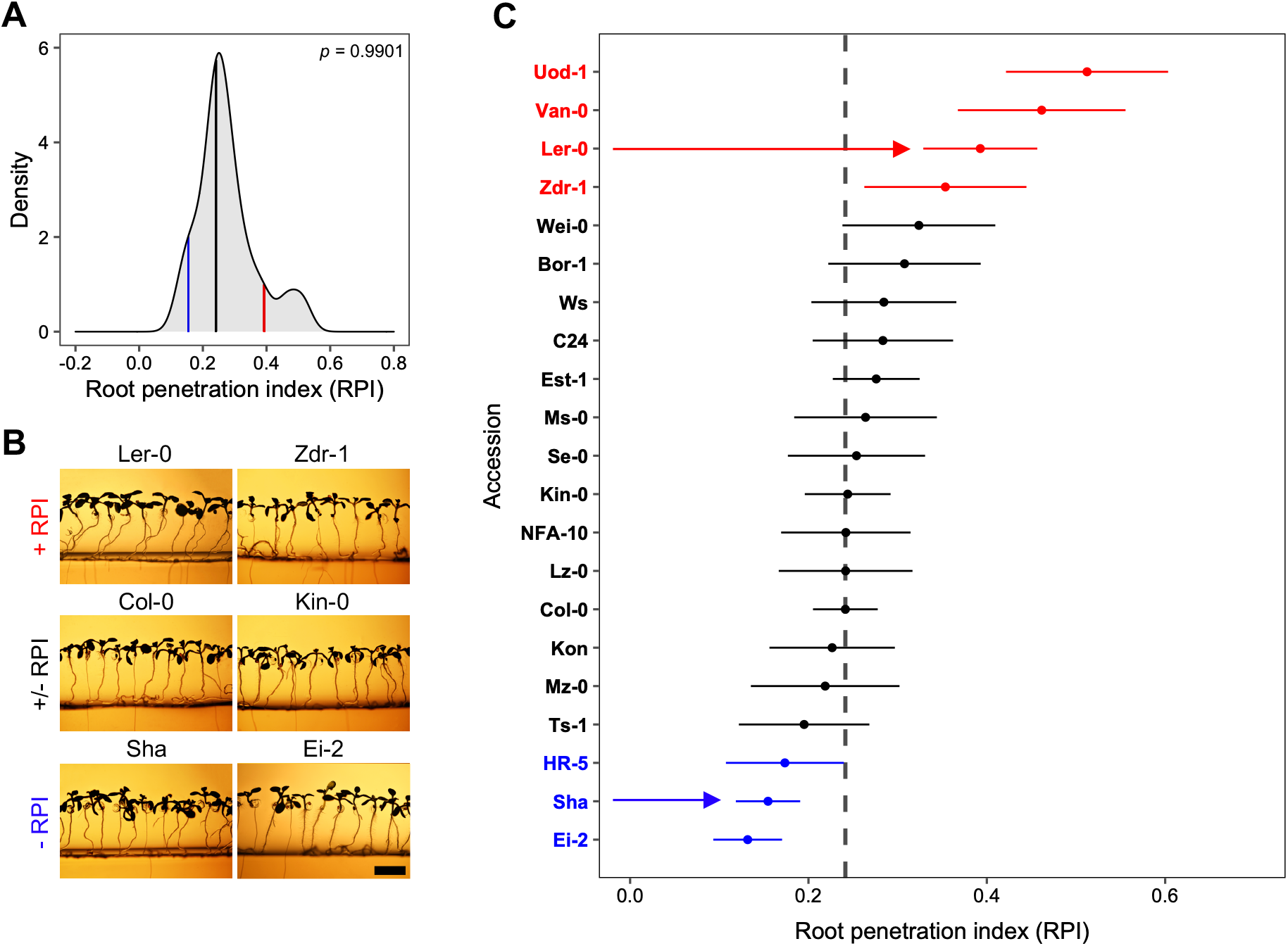
Natural variation in root penetration index (RPI) among Arabidopsis accessions. (A) Density plot of the RPI phenotypes displays a bimodal distribution (Hartigan’s dip test, p<0.05). *p*, p-value; blue line, RPI of Sha; black line, RPI of Col-0; red line, RPI of Ler-0. (B) Representative Arabidopsis accessions that show high, intermediate and low RPI phenotypes at 12 DAS. Scale bars, 4 mm. (C) RPI values of 21 Arabidopsis accessions ranked in ascending order. x axis, RPI trait; y axis, Accessions; colored dots, estimated RPI values; colored lines, confidence intervals; dark grey dashed line, RPI of Col-0. Colors represent different phenotypic groups according to Col-0 accession (chi-square and Fisher’s exact test, p<0.05). In blue, Accessions < Col-0; in black, Accessions = Col-0 and in red, Accessions > Col-0. Data are from three independent experiments (n= 6 plates, with 45 to 50 seedlings per plate).

### q-RPI3 QTL on chromosome 3 governs Arabidopsis root system penetrability

To identify genomic regions associated with root penetration ability, we focused on the accessions Landsberg *erecta* (Ler-0) and Shahdara (Sha), because they have contrasting high and low RPI values, respectively, and because a recombinant inbred line (RIL) population between these two accessions is available. This RIL population has been successfully used to map quantitative trait loci (QTLs) for various abiotic stress tolerance in Arabidopsis (Clerkx *et al*., 2004; Ren *et al*., 2010). A total of 114 RILs were used for phenotyping the RPI trait. For the Ler-0 x Sha mapping population, a large variation in RPI values (ranging from 0.03 to 0.61, Fig. 5A) was observed, suggesting transgressive segregation (Fig. 5B, C, Supplementary Table S5). The observed variation in RPI was under genetic control as indicated by its broad-sense heritability (H^2^ = 0.88). Additionally, frequency distribution of the untransformed RPI data showed a bimodal distribution, suggesting that at least one major QTL was linked to the RPI phenotype (Fig. 5A). Using the constructed genetic map and the phenotypic data obtained from the RIL population, a single QTL mapping was performed. For the RPI trait, a large-effect QTL, hereinafter called *ROOT PENETRATION INDEX 3* (*q-RPI3),* was detected on the top of chromosome 3 (Fig. 6A). *q-RPI3* QTL maps between NT204 and MSAT3.19 markers, showing a highest logarithm of odds (LOD) score around 15 cM near marker MSAT3.19 (Supplementary Fig. S5A). According to this, the *q-RPI3* credible interval comprised a region from 11 to 19 cM, encompassing a 2.48 Mb region on chromosome 3 (1 cM = 311 kb) (Clerkx *et al*., 2004). This genomic region contains 1751 predicted genes (Supplementary Table S6). *q-RPI3* was supported with a LOD score of 8.82 and accounted for 29.98% of phenotypic variance explained (PVE). Further analysis at MSAT3.19 marker indicated that the Ler-0 alleles are positively associated with an increased RPI in *q-RPI3* QTL (Fig. 6B). To assess the presence of additional QTLs and test possible additive and epistatic interactions, we performed a multiple QTL mapping (MQM). The MQM model revealed two additional minor-effect QTLs on chromosomes 2 and 4 (Supplementary Fig. S6). These minor QTLs showed low LOD scores (LOD <3.5) and accounted for less than 8.5% of PVE (Supplementary Fig. S6, Supplementary Table S7). Under MQM assumption, no strong evidence of additive and epistatic interactions was detected (Supplementary Table S7). Based on the foregoing, minor QTLs were not further considered in our analysis. These results provide insights into the genetic architecture behind root penetrability *in A. thaliana* and show that a major QTL (*q-RPI3*) is associated to natural genetic variation in the RPI phenotype among the Ler-0 x Sha RIL population.

**Fig. 5.**
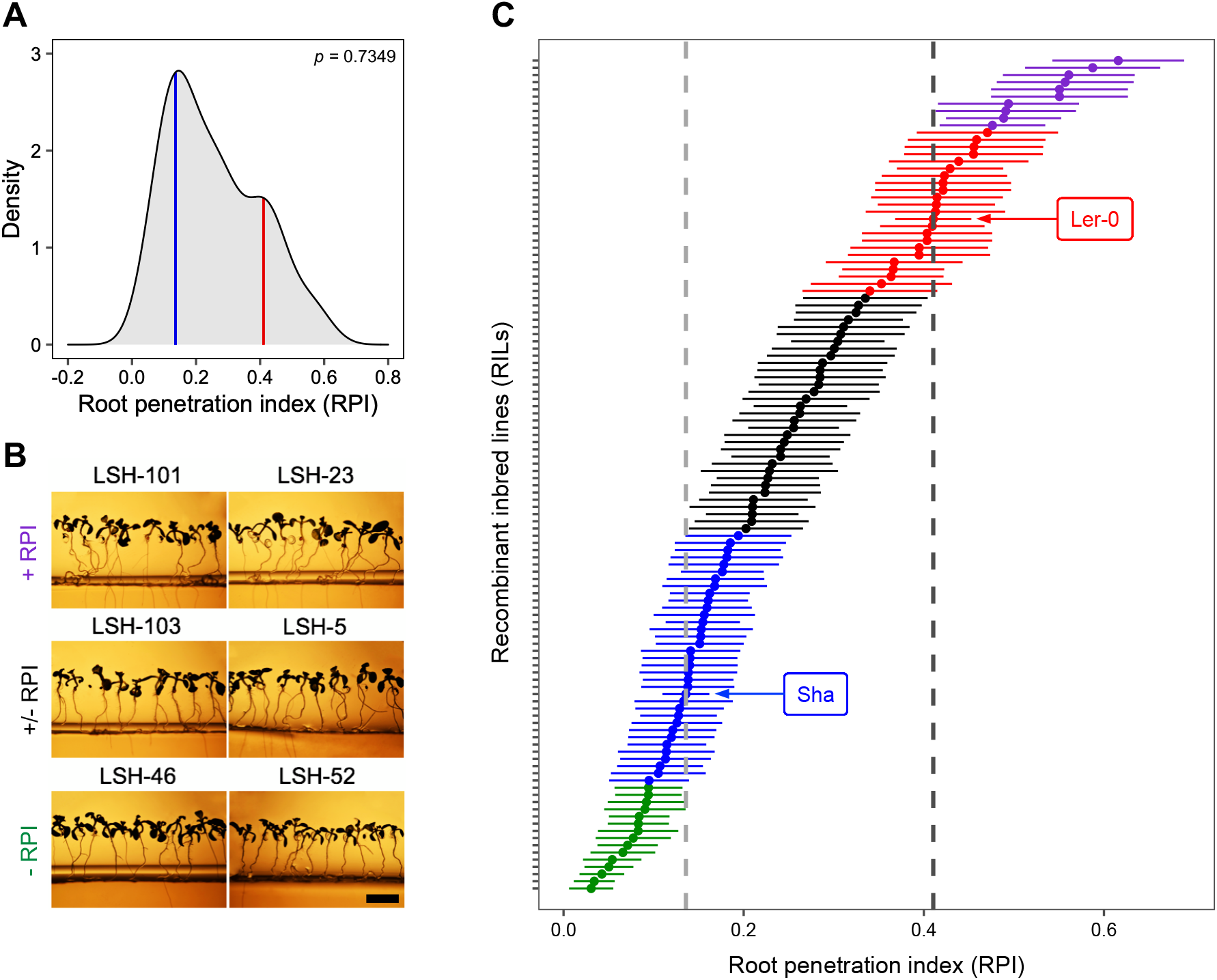
Distribution of the root penetration indexes (RPIs) in the Ler-0 x Sha population. (A) Density plot of the RPI phenotypes displays a bimodal distribution (Hartigan’s dip test, p<0.05). *p*, p-value; blue line, RPI of Sha; red line, RPI of Ler-0. (B) Representative recombinant inbred lines (RILs) that show high, intermediate and low RPI phenotypes at 12 DAS. Scale bars, 4 mm. (C) Root penetration indexes (RPIs) of 114 RILs and parental genotypes ranked in ascending order. x axis, RPI trait; y axis, RILs; colored dots, estimated RPI values; colored lines, confidence intervals; light grey dashed line, RPI of Sha; dark grey dashed line, RPI of Ler-0. Colors represent phenotypic groups with significant differences in reference to parental accessions (chi-square and Fisher’s exact test, p<0.05). In green, RILs < Sha; in blue, RILs = Sha; in black, Ler-0 > RILs > Sha; in red, RILs = Ler-0 and in purple, RILs > Ler-0. Data are from three independent experiments (n= 3 plates, with 45 to 50 seedlings per plate)

**Fig. 6.**
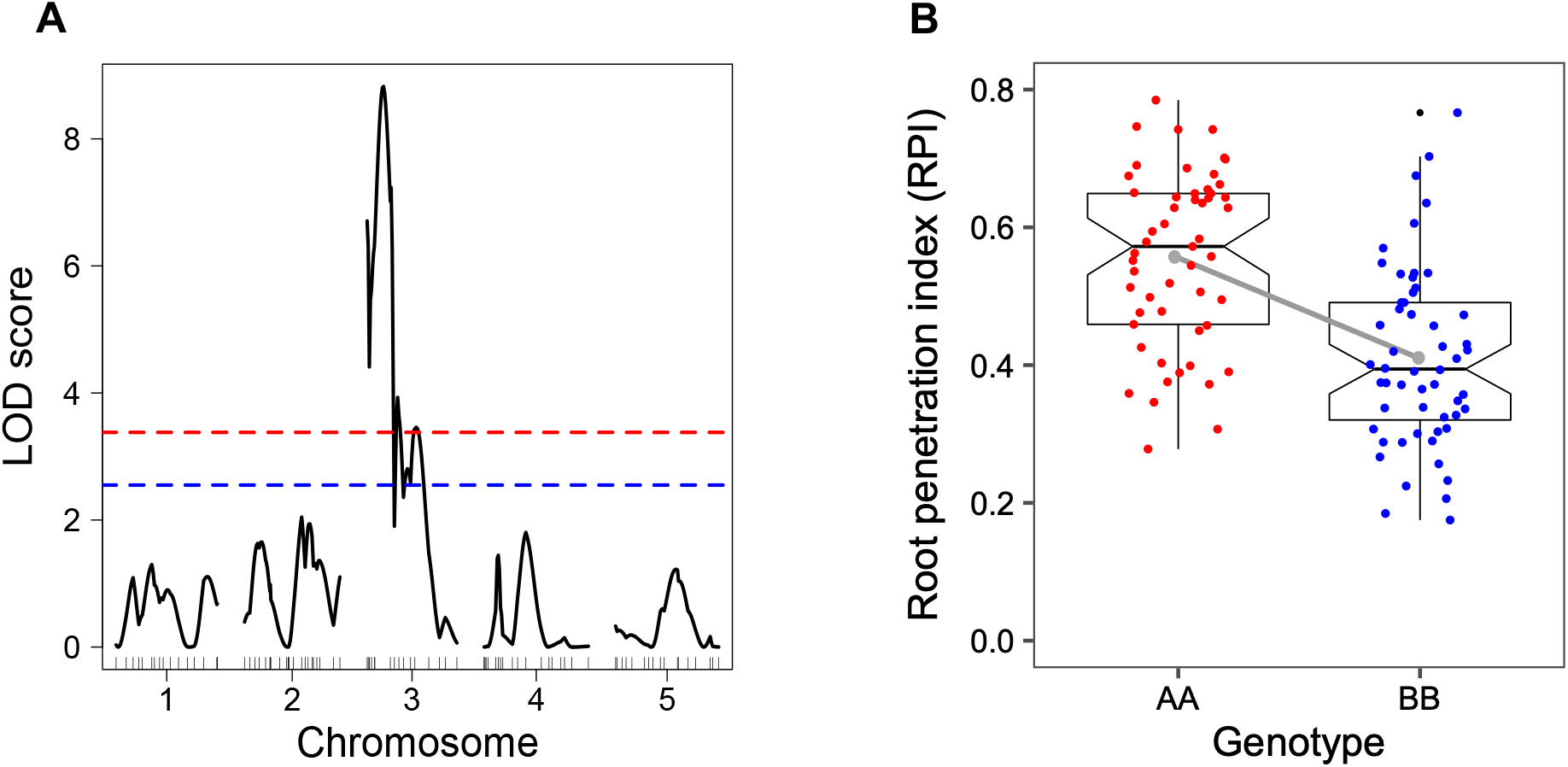
Single QTL analysis for root penetration index (RPI) in the Ler-0 × Sha RIL population. (A) Identification of a major QTL for RPI phenotype on chromosome 3 (*RPI3*). Red and blue dashed lines; significant LOD thresholds (p=0.01 and p=0.05, respectively) using 1000 permutations. (B) Plot of the QTL effect at MSAT3.19 marker. Red and blue dots, RPI values grouped by genotype. Gray line marks the RPI averages among the genotypic groups. AA, Ler-0 allele; BB, Sha allele.

As Sha alleles were negatively connected to root system penetrability, we decided to investigate the genomic collinearity between Ler-0 and Sha accessions. A 2.48 Mb inversion (ranging from ∼2.8-5.3 Mb) on the top arm of chromosome 3 of the Sha genome was previously identified (Jiao and Schneeberger, 2020). A detailed genomic analysis indicated that this chromosomal inversion is localized inside the confidence intervals (∼3.4-5.9 Mb) of the *q-RPI3* QTL (Supplementary Fig. S5B), suggesting that such Sha inversion could be tightly associated with root penetration phenotype in Arabidopsis.

### IL-321 introgression line exhibits a decreased root penetration ability into hard gels

To further validate the relationship between *q-RPI3* and Sha inversion identified on chromosome 3, we evaluated the root penetration ability of the IL-321 introgression line, which carries a homozygous region (∼0.2-5.5 Mb) with the complete inversion and most of the confidence intervals of *q-RPI3* from Sha accession in the Ler-0 genetic background. (Fig. 7A). We observed that the IL-321 root system penetration differs significantly from its parental accessions and shows a significantly lower RPI (RPI=0.28) than the Ler-0 accession (RPI=0.42, Fig. 7B, C), confirming that Sha region is negatively correlated to RPI phenotype. Consequently, these observations confirm that *q-RPI3* contains alleles largely responsible for modulating root system penetrability in these *Arabidopsis* accessions.

**Fig. 7.**
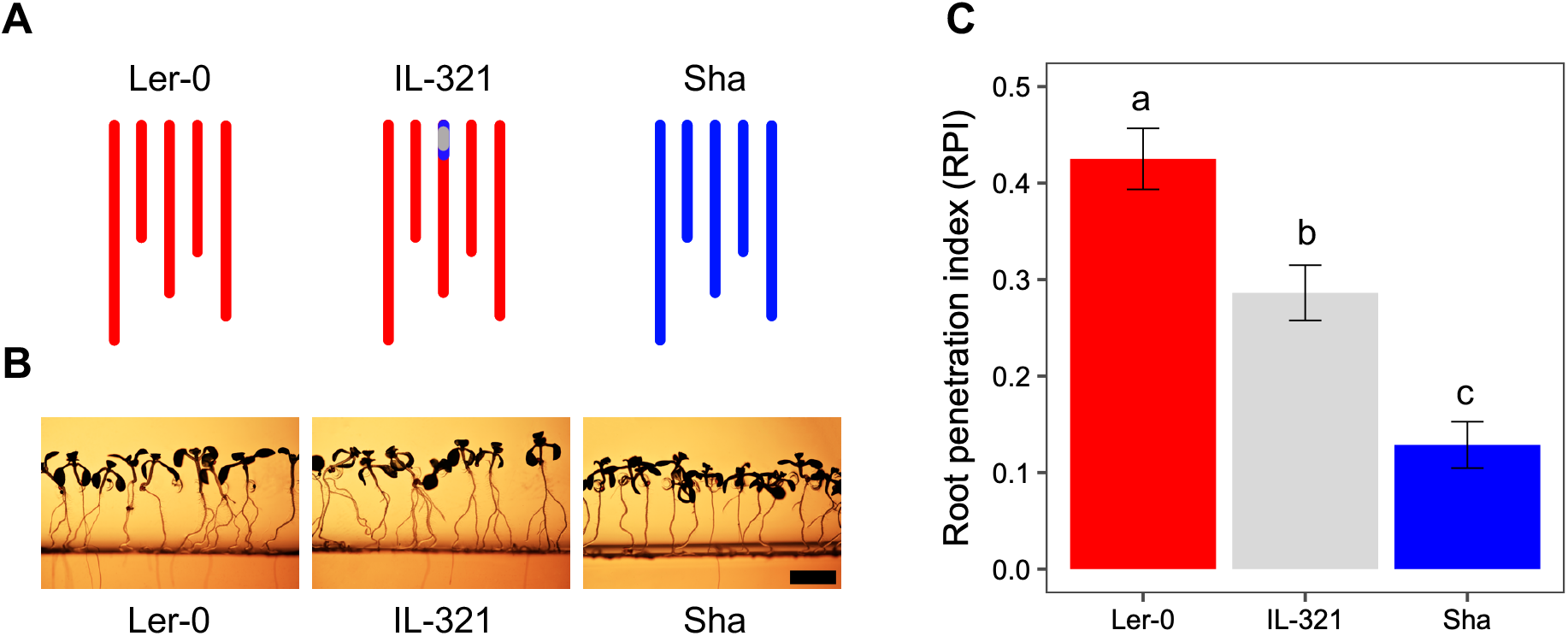
RPI analysis of the IL-321 introgression line (IL) in the two-phase-agar system. (A) Graphic representation of the introgressed region with the chromosomal inversion of Sha in the Ler-0 genetic background. Gray line, chromosomal inversion. (B) Representative images of the IL-321, Ler-0 and Sha accession in the two-phase-agar system at 12 DAS. Scale bars, 4 mm. (C) RPI values of IL-321 and parental genotypes. x axis, lines; y axis, RPI trait; colored bars, estimated RPI values; error bars, confidence intervals; letters, significant differences (chi-square and Fisher’s exact test, p<0.05). Data are from three independent experiments (n= 10-12 plates, with 45 to 50 seedlings per plate).

### NCED3 does not influence the variation in root penetrability in Ler-0 and Sha

Interestingly, *q-RPI3* was colocalized with a previously detected QTL for ABA accumulation in the same Ler-0 x Sha RIL population, where the *9-cis-epoxycarotenoid dioxygenase3* (NCED3) gene, encoding a key enzyme for ABA synthesis, is located (Kalladan *et al*., 2019). Since it has been reported that ABA regulates plant responses to soil compaction (Hartung *et al*., 1994; Tracy *et al*., 2015), we decided to test the possibility of NCED3 being the gene behind the influence of *q-RPI3* on root penetration. Our experiments consisted in assessing primary root penetrability of *nced3* seedlings compared to the WT, and to test the effect of ABA on root penetrability in Ler-0, Col-0, Sha and IL-321. In the latter case, we tested the effect of different ABA concentrations (0, 0.1, 1 and 3 µM) on root penetrability. We did not find any significant difference in root penetration ability between Col-0 and *nced3* mutant (Supplementary Fig. S7A). This result suggests that, under our experimental conditions, the NCED3-induced ABA accumulation is probably not responsible for the differences in root penetrability between Ler-0 and Sha. Exogenous ABA application on Ler-0 and IL-321 line had a negative impact on in their root penetrability, whereas it did not have effect on Col-0 and Sha. These observations suggest that ABA does not play a major role on primary root penetrability (Supplementary Fig. S7B). Further analyses to fine mapping and discovery of candidate genes in the *q-RPI3* QTL interval are needed.

## Discussion

In the present study, we performed an analysis of the architectural plasticity during Arabidopsis root system penetration into agar layers with low and high mechanical impedance. During the standardization of the two-phase-agar system, we observed strong correlations between agar concentration and the studied variables: PMR, RPI and RD (Fig. 1, Supplementary Table S1). Diverse studies have demonstrated that the gelling agents (phytagel or agar) at different concentrations can modify the mechanical properties of culture media, promoting sundry root growth patterns such as root waving (Okada and Shimura, 1990), skewing (Yamamoto *et al*., 2008) and bending (Yan *et al*., 2018). Our results show that higher concentrations of agar (higher mechanical impedance) drastically restrict both primary root penetration and rooting depth (Fig. 1, Supplementary Table S1). Similar results were obtained by employing phytagel media to mechanically impede Arabidopsis roots, where increasing phytagel concentrations negatively correlated with root length and root penetration percentage (Yan *et al*., 2017). Such root responses could be related to a decrease in the number and size of pores present on the surface of media containing a higher gel strength (Narayanan *et al*., 2006) or a higher mechanical resistance of the gel to be penetrated by roots. By using specific agar concentrations in the growth medium, it is feasible to analyze root-gel interactions, including root responses to mechanical impedance and natural variation in root penetration ability.

As a proof of concept, we established an experimental design to evaluate the Arabidopsis seedling responses to agar mechanical impedance during three root penetration stages. The phenotypic analysis at two agar concentrations indicated that the seedlings which root failed to penetrate the bottom agar layer were severely affected by agar impedance, showing drastic decreases in root penetration, leaf area, root length and size of elongation zone under both mechanical treatments (Fig. 2, Supplementary Fig. S2). Also, a significant increase in root diameter and an ectopic formation of root hairs near root tip were observed (Fig. 2E, K). Similar morphologies have been reported for other plant species exposed to mechanical impedance (Colombi and Walter, 2016; Potocka and Szymanowska, 2018). In general, plants exhibit a marked decline in root system growth and penetration when they are subjected to soil mechanical stress (Tracy *et al*., 2012; Potocka and Szymanowska, 2018). Such root responses seem to be consequence of a complex mechanism that involves the inhibition of root cell elongation mediated by a cross-talk between ethylene and auxin signaling (Santisree *et al*., 2011); Okamoto *et al*., 2021). In parallel, a recent transcriptional profiling of Arabidopsis impeded roots revealed that early root responses to mechanical impedance are specifically orchestrated by a network of ROS, ethylene and auxin signalling (Jacobsen *et al*., 2021). We found that seedlings whose roots were able to penetrate the second agar layer seem to be less affected, except for a slight reduction in root length, depth and diameter, than those that failed to penetrate (Fig. 2, Supplementary Fig. S2). These results suggest that once the primary root penetrates a higher impedance barrier, normal seedling growth resumes. A recent publication reported that highly-penetrating maize genotypes into compacted soil, showed 50% greater dry shoot biomass than genotypes with poor root penetration (Schneider *et al*., 2021). Our results are in line with previous reports suggesting that deep root penetration into soil horizons furnishes diverse adaptive advantages for crops to grow in harsh environments (Uga *et al*., 2013. This makes the study of the genetic and molecular mechanisms controlling root system plasticity under soil compaction conditions a primary topic of interest in plant biology.

To dissect the molecular responses that allow Arabidopsis primary root penetration in agar layers, we studied the functional role of auxins during root morphological responses to continuous mechanical impedance. In mechanically obstructed roots, we observed an increased *DR5::GFP* signal, expanding from the root tip to the elongation zone (Fig. 3A, C). The phenotypes of the roots unable to penetrate, including a reduction in root elongation, a waving growth pattern and an ectopic induction of root hairs near the root tip, could be correlated to a mechanical stress-induced auxin transport or activity. In addition, the spatio-temporal expression of the PIN2 auxin efflux carrier detected in the elongation zone of the roots (in both mechanical impedance conditions) revealed an interesting expression pattern for each of the different penetration stages, indicating that the dynamics of auxin distribution in root tips may also plays a key role during the penetration process into hard agar layers. Many studies have shown that root morphology alterations under high mechanical impedance are triggered by an ethylene-promoted auxin biosynthesis (Okamoto *et al*., 2008; Okamoto *et al*., 2021). Such idea is consistent with the current model of ethylene-induced auxin biosynthesis in the root meristem and its transport to the elongation zone by different auxin efflux carriers (PIN family and AUX1). Then, auxin accumulation in root-specific tissues inhibits cell elongation and elicits changes in root growth patterns (Ruzicka *et al*., 2007). In agreement with ethylene and auxin-mediated root growth inhibition, it has also been reported that in tomato ethylene-induced auxin synthesis and/or transport play a role in determining root penetration capacity (Santisree, *et al*., 2011). Altogether, the *DR5::GFP* and *PIN2::PIN2::GFP* expression patterns indicate that, similar to bending responses observed in root thigmotropism, gravitropism and halotropism, PIN2-medited active redistribution of auxin plays a critical role during the root growth responses to mechanical impedance (Galvan-Ampudia *et al*., 2013; Sato *et al*., 2015; Lee *et al*., 2020). In general, our findings suggest that auxin biosynthesis and redistribution (across specific cell types and root zones) are two physiologically important processes associated to different stages of root penetration into agar media.

Along with the molecular mechanisms, it also important to decode the genetic basis behind Arabidopsis root system penetrability. Using a small set of accessions from highly diverse edaphic environments, we showed that there is a considerable natural variation in the capacity of Arabidopsis root system to penetrate hard agar layers (Fig. 4A, Supplementary Table S4). As shown in Fig. 4C, we observed that the root penetrance of Arabidopsis accessions presents a continuous variation from a minimum RPI of 0.13 for the Ei-2 accession to a maximum RPI of 0.51 for the Uod-1 accession. Several studies have documented the natural variation in root penetrability in monocot and dicot plant species (Yu *et al*., 1995; Bushamuka and Zobel, 1998; Klueva *et al*., 2000; Whalley *et al*., 2012). It has been proposed that differences in root diameter *per se* and maximum root growth pressure were responsible of the root penetrability variation among plant species (Materechera *et al*., 1991). However, recent reports on maize, wheat and barley showed that such variation is more strongly associated with variation in specific root anatomical traits that are negatively regulated by ethylene. This suggest that plant root systems with high ethylene insensitivity could have major advantages of growing in compact soils (Pandey *et al*., 2021; Schneider *et al*., 2021; Vanhees *et al*., 2021). Despite these advances, further analyses are still required to establish whether natural variation in root system penetration is also controlled by ethylene-induced anatomical changes in dicot roots. Also, it remains unclear how natural genetic variability mediates root penetration in response to soil mechanical constraints and the role of plant hormones other than auxin and ethylene play in this variation.

Considering the natural genetic variation observed, we performed a QTL mapping in a Ler-0 x Sha RIL population that allowed the identification of a QTL associated to primary root penetrability in chromosome 3 of Arabidopsis (Fig. 6A). *q-RPI3* QTL accounted for 29.98% of the total PVE in the RILs under study, suggesting that *q-RPI3* underlies a significant proportion of the RPI. In addition, the *q-RPI3* candidate genomic region overlaps with a previous identified 2.48Mb inversion in chromosome 3 of Sha accession, calling into question the functional role that such inversion in root penetration (Supplementary Fig. S5B) (Jiao and Schneeberger, 2020). By phenotyping primary root penetrability in the introgression line IL-321, we validated the contribution of *q-RPI3* in modulating the natural variation in the capability of Arabidopsis roots to penetrate strong agar layers (Fig. 7C). The lower primary root penetrability observed in IL-321 agrees with a negative effect of Sha alleles. Several QTLs for root penetration ability have been detected in monocot plants, such as rice (*Oryza sativa* L.) (Ray *et al*., 1996; Zheng *et al*., 2000; Zhang *et al*., 2001) and wheat (*Triticum* spp.) (Kubo *et al*., 2007; Botwright Acuña *et al*., 2014). For instance, QTL studies in rice identified a total of 35 QTLs associated with the RPI trait, which account for a phenotypic variation ranging from 3.84% (small effect QTL) to 26.2% (large effect QTL) (Courtois *et al*., 2009). Interestingly, a genomic region that covers 15 Mb (20-35 Mb) groups 8 QTLs from different mapping populations, suggesting a stable genetic component in common among mapping populations (Courtois *et al*., 2009). On this basis, many recombinants inbred near-isogenic lines (RINILs) have been generated (Clark *et al*., 2008). The analysis of such RINILs revealed that root penetration ability is associated with greater bending stiffness in superior genotypes (RINILs 9 and 11) (Clark *et al*., 2008). Likewise, UDP-glucose 4 epimerase 1 (OsUGE-1) on rice chromosome 7 has been suggested as a candidate QTL/gene responsible for modulating the thickness of penetrating roots and closely linked to root penetration into wax-petrolatum layers (Nguyen *et al*., 2004). However, until now the direct relationship between OsUGE-1 and root penetration ability remains to be investigated. Other promising candidate genes are Zm00008a033967 (MEI2-like RNA binding protein) that is involved in the formation of multiseriate cortical sclerenchyma in maize, and histidine kinase-1 (HK1) that modulates root circumnutation in rice; both genes are presumably associated to root penetration capacity into compact substrates (Schneider *et al*., 2021; Taylor *et al*., 2021). Finally, we did not find any strong evidence of the involvement of NCED3 and ABA between the Ler-0 and Sha accessions. Further investigations are required to evaluate other candidate genes in *q-RPI3* QTL interval and to more carefully test the involvement of ABA on root penetrability. The present study demonstrated that *q-RPI3* is crucially important for a high primary root penetration in *A. thaliana*. Identifying the genes behind the role of *q-RPI3* on root penetration is an on-going task to gain a better understanding on the genetic mechanism allowing plants to tolerate diverse soil adversities. Due to considerable importance of root penetrability in contributing to crop productivity under soil compaction conditions, it is necessary to advocate for novel plant breeding strategies to develop soil compaction-resilient crop varieties.

## Abbreviations

ABA: abscisic acid
IR: impeded roots
MCA1: *mid1*- complementing activity 1
NCED3: *9-cis-epoxycarotenoid dioxygenase3*
NDR: non-disturbed roots
PR: penetrating roots
PMR: penetrometer resistance
PRP: primary root penetrability
QTLs: quantitative trait loci
RIL: recombinant inbred lines
RPI: root penetration index
q-RPI3: ROOT PENETRATION INDEX 3
RPS: root penetration stages
RD: rooting depth
SBAL: surface of the bottom agar layer

## Supplementary data

Supplementary data are available at JXB online.

Fig. S1. The *in vitro* system used in this study.

Fig. S2. Morphology of Arabidopsis seedlings grown at three penetration stages under low and high mechanical impedance.

Fig. S3. Morphological characterization of Arabidopsis root system responses to the low and high mechanical impedance.

Fig. S4. Analysis of the *QC46::GUS* and *CycB1;1::GFP* reporter lines during Arabidopsis root penetration into agar layers.

Fig. S5. Major QTL for the RPI phenotype identified on chromosome 3 (*q-RPI3*).

Fig. S6. Multiple QTL mapping (MQM) for the RPI phenotype in the Ler-0 × Sha RIL population.

Fig. S7. Analysis of primary root penetrability among different Arabidopsis lines under the two-phase-agar system.

Table S1. Pearson and Spearmańs correlation coefficients between agar concentration and tested traits.

Table S2. Summary of the tested traits analyzed in the two-phase agar system.

Table S3. Arabidopsis accessions, soil units and physical properties.

Table S4. RPI values of Arabidopsis accessions obtained from the two-phase-agar system.

Table S5. RPI values of the Ler-0 x Sha RIL population obtained from the two-phase-agar system.

Table S6. Number of predicted genes in the *q-RPI3* QTL support interval (95% Bayesian interval) and in the genomic inversion located on the top of chromosome 3.

Table S7. Summary of QTL analysis using a multiple QTL model (MQM) for RPI phenotype in the Ler-0 × Sha RIL population.

Video S1. Time-lapse video of Arabidopsis seedlings growing under low and high mechanical impedance.

## Acknowledgements

We would like to thank Dr. Carlos Alonso Blanco from the Centro Nacional de Biotecnología, Consejo Superior de Investigaciones Científicas, Madrid, Spain, for providing us IL-321 seeds. We thank Dr. Octavio Martínez for his support for statistical analysis. We also thank Araceli Oropeza Aburto for technical assistance.

## Author contributions

EBB and LHE designed research. EBB and TYRC performed experiments. EBB, RRA and LHE analyzed data. RRA and LHE provided reagents and analytic tools. EBB and LHE wrote the paper. All authors have read and approved the final manuscript.

