## Supplementary Figures S1-S7 for "ROOT PENETRATION INDEX 3, a major quantitative trait locus (QTL) associated with root system penetrability in Arabidopsis"

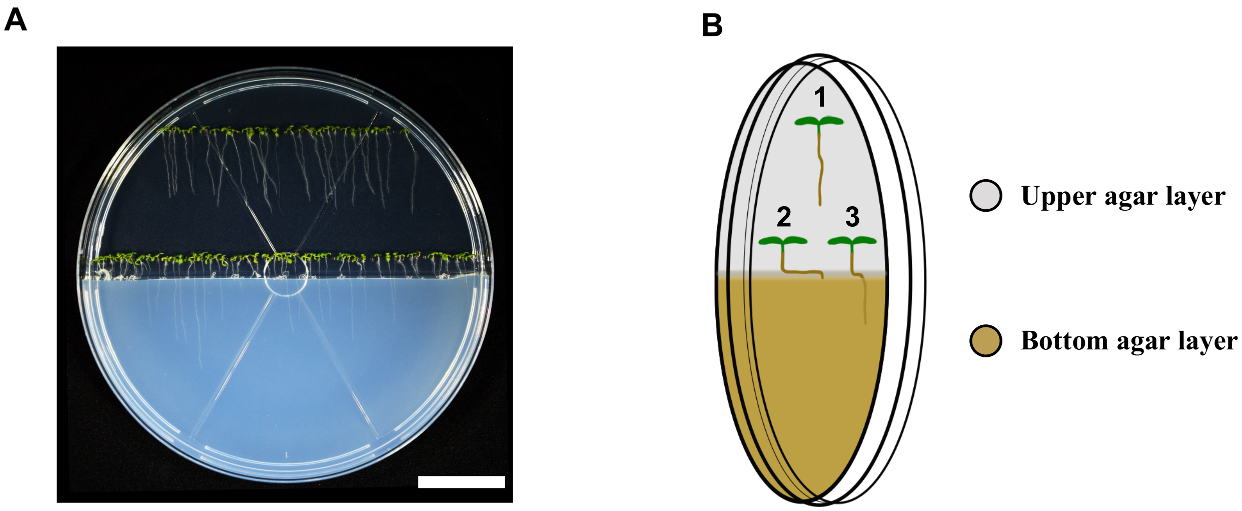


**Supplementary Figure 1. The *in vitro* system used in this study.** (A) Representative image of a petri dish containing between 45 to 50 Arabidopsis seedlings (Col-0) grown under the two-phase-agar system. Scale bar, 3 cm. (B) Graphical representation of the two-phase-agar system. Numbers represent the three root penetration stages (RPS): before (1), during (2) and after (3) Arabidopsis root tips touch the bottom agar layer at low or high mechanical impedance.

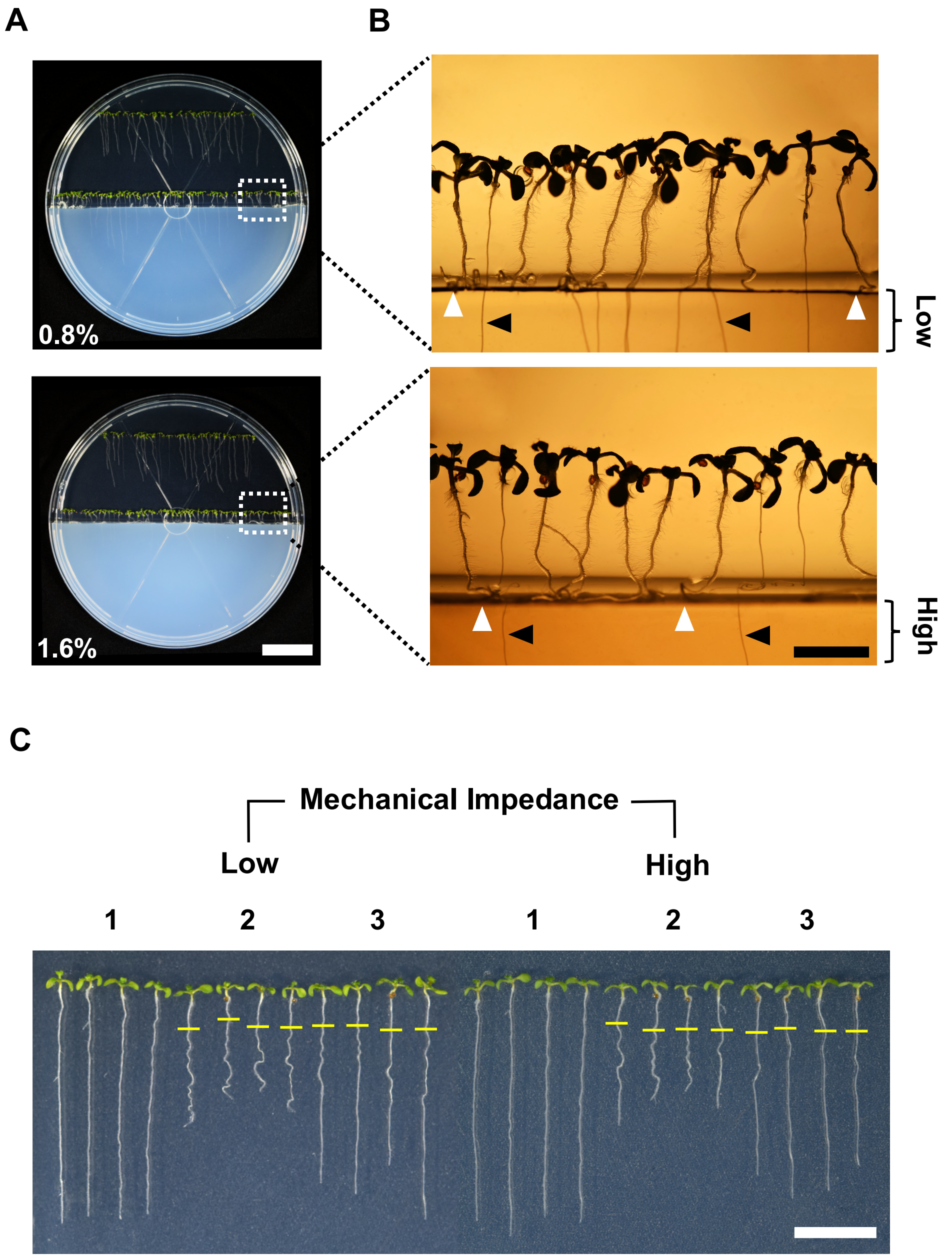


**Supplementary Figure 2. Morphology of Arabidopsis seedlings grown at three penetration stages under low and high mechanical impedance.** (A) Low and high mechanical impedance treatments in the two-phase-agar system (0.8% and 1.6% agar concentration, respectively). Scale bar, 3 cm. (B) Close-up views of Arabidopsis root phenotypes at low and high mechanical impedance. Black arrows, penetrating roots (PR); white arrows, impeded roots (IR). Scale bar, 4 mm. (C) Morphology of 12-day-old Arabidopsis seedlings grown under low and high mechanical stress conditions at three RPS. Seedlings images were fused from two adjacent stacks. Yellow bars, first root-agar contact. Scale bar, 1 cm.

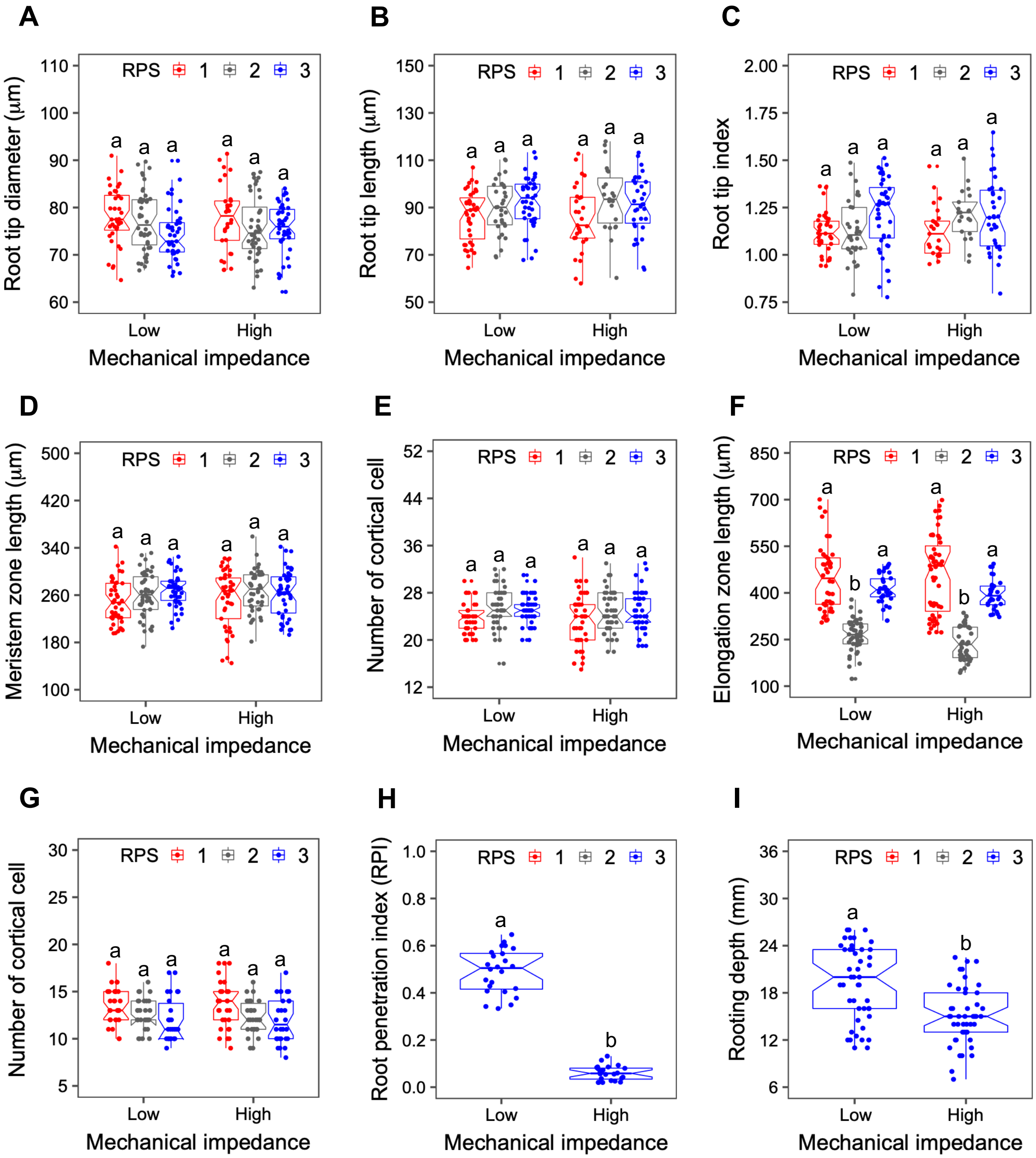


**Supplementary Figure 3. Morphological characterization of Arabidopsis root system responses to the low and high mechanical impedance.** (A-I) Arabidopsis root traits evaluated in three RPS at low and high mechanical impedance. Box plots show the tested morphological traits. Horizontal lines, medians; box limits, 25th and 75th percentiles; letters, significant differences (p=0.01, one and two-way ANOVA and Tukey´s HSD). Data are from three independent experiments (n= 10-12 seedlings per experiment).

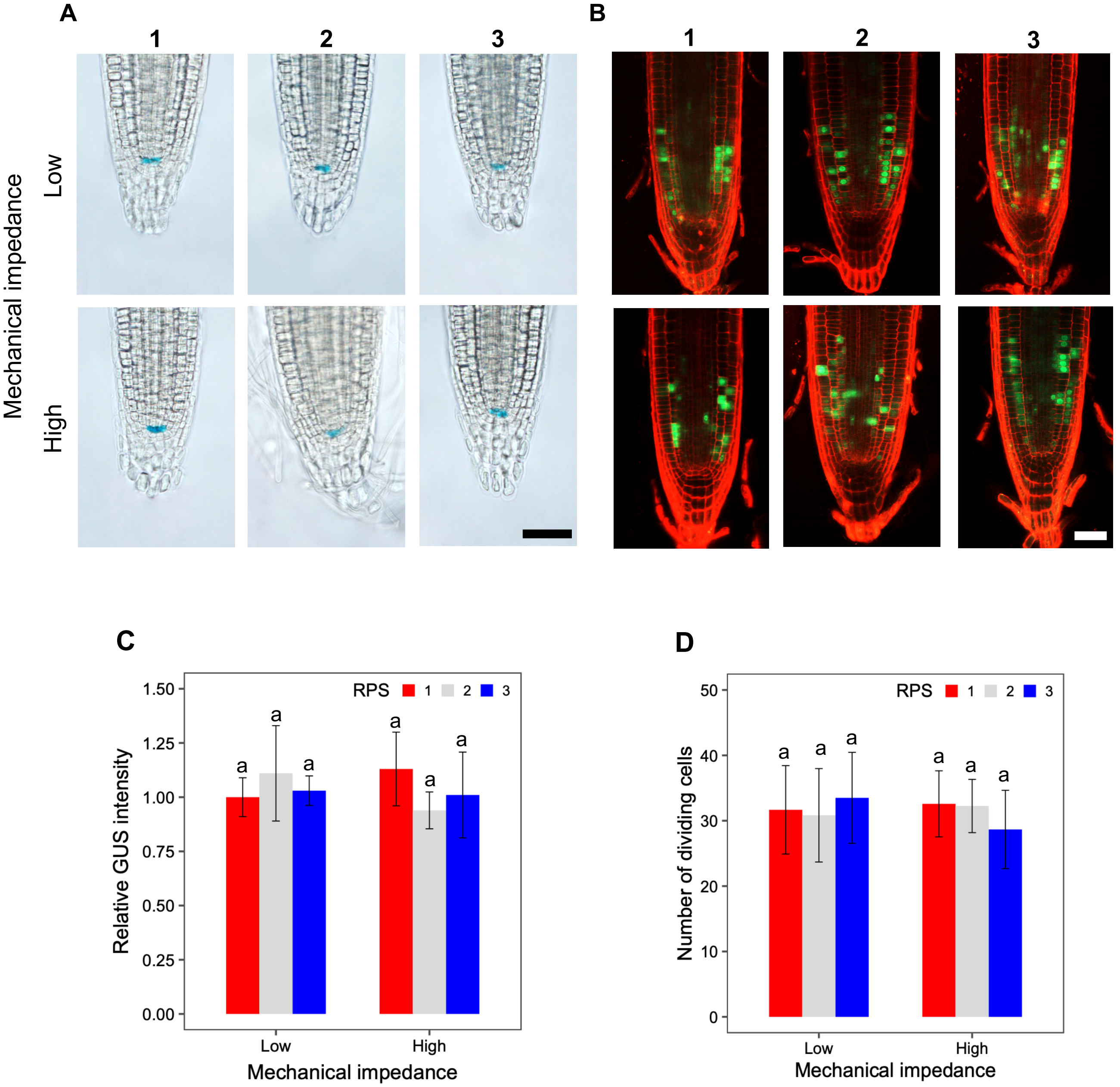


**Supplementary Figure 4. Analysis of the *QC46::GUS* and *CycB1;1::GFP* reporter lines during Arabidopsis root penetration into agar layers.** (A) Expression of *QC46::GUS* in Arabidopsis roots during three RPS at low and high mechanical impedance. Scale bar, 50 µm. (B) Confocal images of the mitotic cyclin CycB1 in Arabidopsis roots grown under two mechanical treatments. Green color, expression of green fluorescent protein (GFP); red color, propidium iodide staining. Scale bar, 50 µm. (C) Relative GUS intensity of the *QC46::GUS* reporter line. (D) Quantification of *CycB1;1::GFP* signal in Arabidopsis roots. Bar plots show the mean ± standard deviation of GUS intensity or GFP fluorescence. Letters, significant differences ($p=0.01$, two-way ANOVA and Tukey´s HSD). Data are from three independent experiments (n= 10-15 roots).

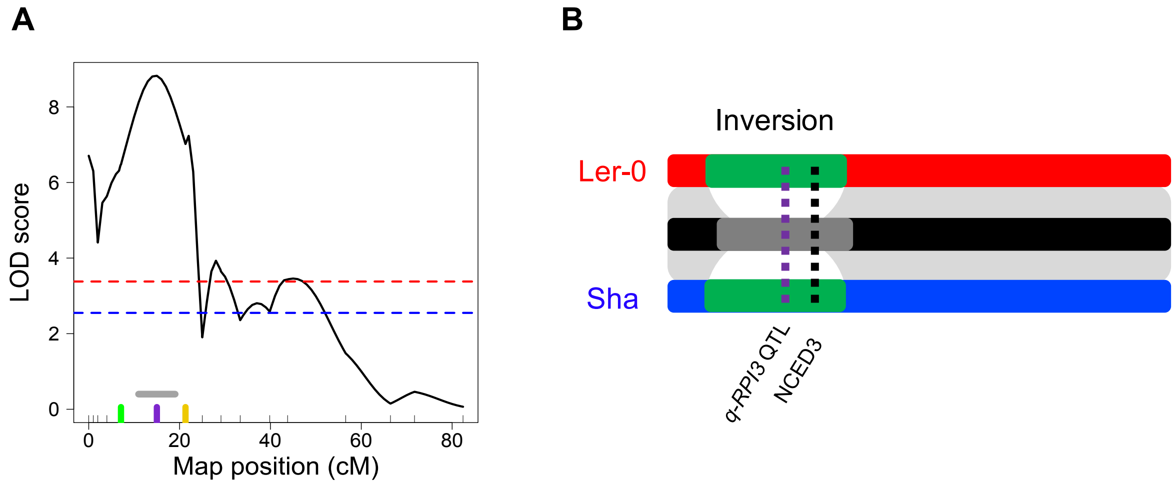


**Supplementary Figure 5. Major QTL for the RPI phenotype identified on chromosome 3 (*q-RPI3*).** (A) *q-RPI3* peak represents the maximum LOD score (LOD = 8.82). Red and blue dashed lines; significant LOD thresholds (p=0.01 and p=0.05, respectively); gray line, QTL support interval (~11-19 cM); green line; NT204 marker; purple line, maximum LOD score (15 cM); yellow line, MSAT3.19 marker. (B) Graphic representation of the co-localization of the chromosomal inversion of Shahdara, *q-RPI3* and NCD3 gene on chromosome 3. Green rectangle, inversion; gray rectangle, QTL interval; purple dashed line, *q-RPI3* localization; black dashed line; NCED3 localization.

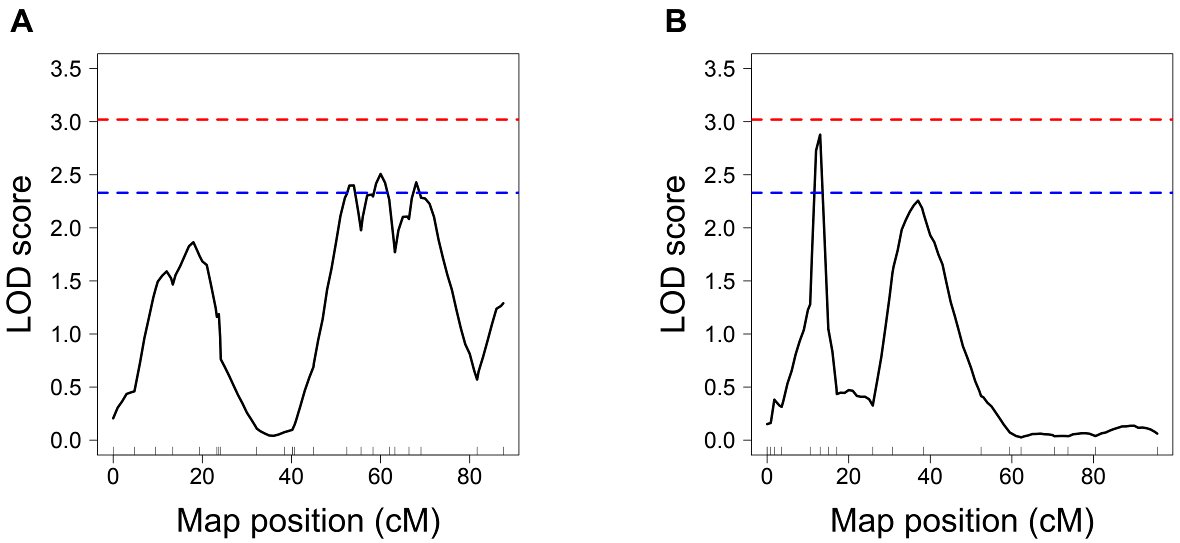


**Supplementary Figure 6. Multiple QTL mapping (MQM) for the RPI phenotype in the Ler-0 × Sha RIL population.** (A) Minor-effect quantitative trait locus (QTL) localized on chromosome 2. (B) Minor-effect QTL identified on chromosome 4. Red and blue dashed lines; significant LOD thresholds (p=0.01 and p=0.05, respectively) using 1000 permutations.

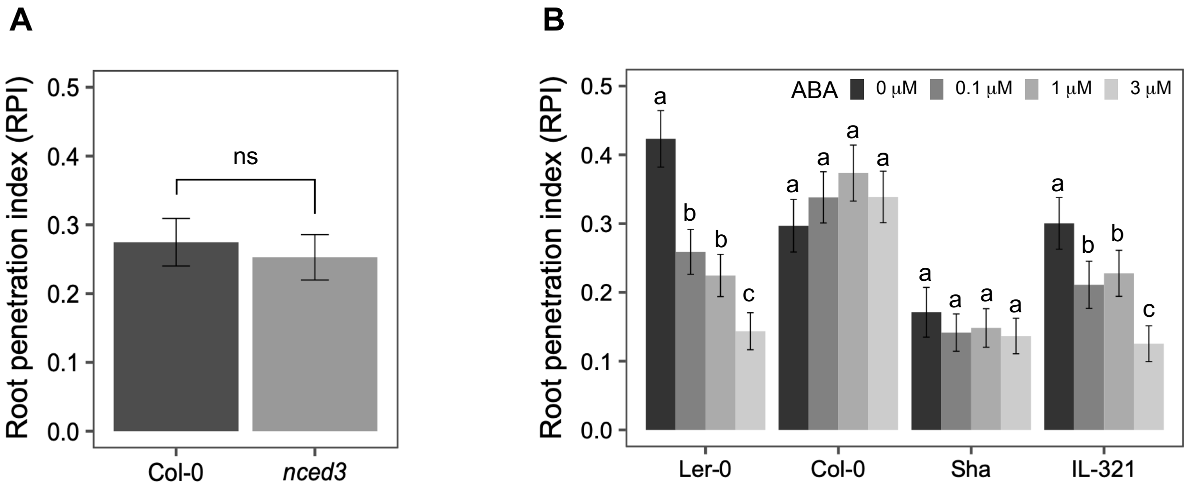


**Supplementary Figure 7. Analysis of primary root penetrability among different Arabidopsis lines under the two-phase-agar system.** (A) Comparative RPI analysis between Col-0 accession and *ncde3* mutant. (B) Effect of exogenous ABA application on root system penetrability of Ler-0, Col-0, Sha and IL-321. x axis, tested lines; y axis, RPI trait; grey bars, estimated RPI values; error bars, confidence intervals; letters, significant differences; ns, not significant; ABA, ABA concentrations (chi-square and Fisher's exact test, $p<0.05$). Data are from three independent experiments (n= 10-12 plates, with 45 to 50 seedlings per plate).
